# Simultaneous ligand binding to intact and partially formed ATP binding sites in the hexameric termination factor Rho

**DOI:** 10.1101/2025.07.22.665897

**Authors:** Tyler D Billings, Kristie Baker, Philip Lacey, Rodrigo Muzquiz, Vicki H. Wysocki, Mark P. Foster

**Author notes:** **Corresponding Author** Mark P. Foster, Email address, Mailing address: 734 Riffe Building, 496 West 12^th^ Avenue, Columbus, Ohio 43210. **Author Contributions** **Tyler D Billings**: Writing, Conceptualization, Resources, Methodology, Formal Analysis, Visualization. **Kristie Baker**: Writing, Investigation. **Philip Lacey**: Writing, Investigation, Formal Analysis, Visualization. **Rodrigo Muzquiz**: Conceptualization, Investigation, Methodology. **Vicki H. Wysocki**: Writing, Resources, Supervision, Funding Acquisition. **Mark P. Foster**: Writing, Conceptualization, Formal Analysis, Supervision, Funding Acquisition.

## Abstract

Thermodynamic coupling between ligand binding sites affords macromolecular machines a means to coordinate processive function. Because these machines may be compositionally complex, quantifying and interpreting ligand binding events can be experimentally difficult. Biophysical methods that convolve binding events into a one-dimensional metric, which suffice for monomeric macromolecules that bind to a single ligand, are insufficient to adequately describe the complexity of binding to oligomeric systems. Confounding factors include structural heterogeneity that may invalidate basic assumptions used to interpret the measurements. In this communication, we use native mass spectrometry to measure ATP binding to a hexameric helicase, the *E. coli* termination factor Rho. Providing new insights into classical and more recent biochemical experiments, we observe and quantify ATP binding to hexameric and lower-order complexes. Moreover, we observe super-stoichiometric binding consistent with ATP binding to partially formed binding sites at the edges of the open washer structure. Such detailed insights are likely critical to understanding the mechanisms by which a broad range of macromolecular machines harness the free energy from ligand binding, hydrolysis, and exchange to coordinate their ligand-dependent functions.

## Introduction

Mechanistic insights into ligand-altered protein function frequently depend on detailed thermodynamic parameters describing free energy changes associated with ligand binding and coupling to allosteric sites.(1–5) Thermodynamic parameters are conventionally obtained via ligand titrations followed by fitting a concentration-dependent signal to a suitable binding model. Such approaches are generally robust when the target protein is monomeric and binds to a single ligand. However, in the case of oligomeric proteins, proportionality between modeled populations and the measured signal may be lost.(6) A major confounding factor is that the presence of multiple oligomeric states can result in signals that reflect the superposition of multiple distinct isotherms, requiring additional experiments to account for the various species and the equilibria that link them.(7, 8)

Hexameric helicases are a class of motor proteins for which detailed ligand binding thermodynamic parameters can be important for understanding their function. These enzymes translocate unidirectionally on single-stranded nucleic acids by converting the chemical potential of nucleotide triphosphates into motion. Several hydrolytic mechanisms have been proposed, with the rotary model of NTP hydrolysis emerging as well suited to explain rapid, directional translocation along polynucleotide substrates.(9) Experimental quantification of the thermodynamic coupling between hydrolytic sites could provide strong evidence in support of the coordinated and sequential ligand exchange underlying the rotary mechanism. Such coupling terms can be extracted by measuring populations of liganded states over a suitable concentration range and fitting with an appropriate mechanistic model that incorporates these values.(2, 5)

A factor complicating the recording and analysis of ligand titration data for this class of enzyme are heterogeneous oligomeric populations under biochemically-relevant concentrations.(10–20) A prominent example from the hexameric helicase superfamily is *E. coli* Rho (Rho, Figure 1A).(21) Conventional ligand binding measurements exhibit multiphasic isotherms, which have been explained by the presence of three weak and three strong ATP binding sites, with dissociation constants in the range of 10 and 100 µM, respectively.(22–24) A potentially confounding factor in fluorescence-based binding assays is that allosteric effects can alter the observed fluorescence signal, which is generally assumed to be proportional to ligand binding to the hexameric form of Rho.(11)

**Figure 1.**
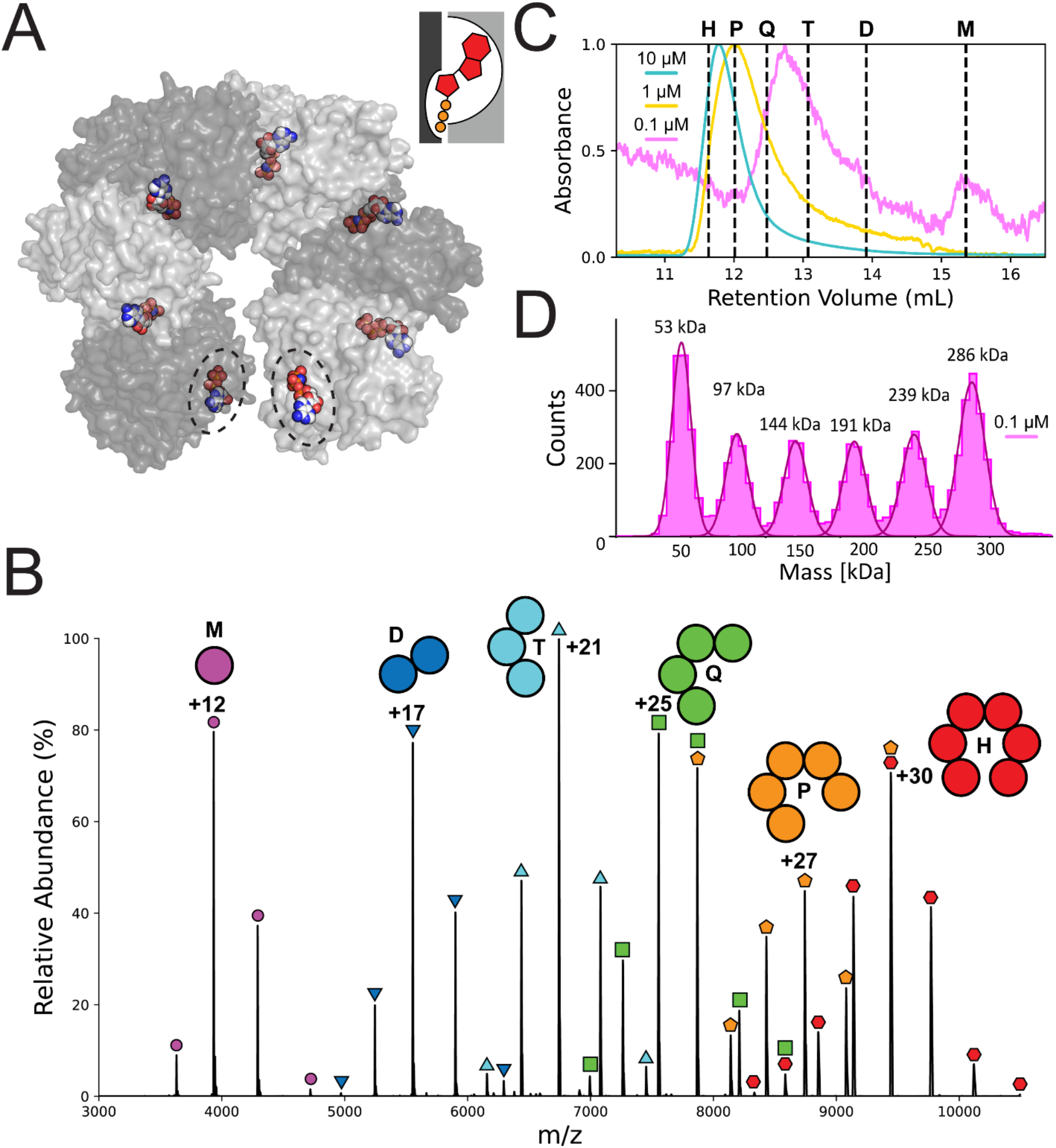
Rho adopts multiple oligomeric states observable in solution and in the gas phase. A) Molecular surface representation of Rho as crystallized in an open-ring hexameric nucleotide-bound form (PDB 1PVO).(21) A complete ATP binding site consists of one subunit contributing 80% of the buried ligand surface area, and the adjacent subunit the remaining 20%, illustrated by the inset diagram. Partially formed binding sites at the edge of the open-ring hexamer are circled. B) Native mass spectrometry of Rho recorded at 10 µM protomer concentration reveals a series of charge states corresponding to the entire range of oligomeric species, including monomer (circles, 47,198.40 Da predicted, 47,196.34 ± 0.22 Da observed), dimer (inverted triangles, 94,396.80 Da predicted, 94,399.42 ± 2.79 Da observed), trimer (triangles, 141,595.20 Da predicted, 141,593.10 ± 4.60 Da observed), tetramer (squares, 188,793.60 Da predicted, 188,823.10 ± 4.41 Da observed), pentamer (pentagons, 235,992 Da predicted, 236,047.20 ± 9.23 Da observed) and hexamer (hexagons, 283,190.40 Da predicted, 283,282.50 ± 20.22 Da observed); the narrow charge distributions are consistent with native-like states for each species. C) Analytical SEC as a function of concentration reveals a shift in the average retention volume of Rho and the emergence of a monomer peak at low Rho protomer concentrations. D) MP reveals a much more complex mixture of populations at 0.1 µM protomer concentration.

Native mass spectrometry (nMS) has the capacity to differentiate between oligomeric species in a heterogeneous sample.(25–32) Measurement of small molecule ligand binding to hetero- and homo-oligomeric protein complexes has also been demonstrated using nMS. (2, 3, 15, 29, 33–35) In this study, we leverage these strengths to measure ATP binding to Rho, which populates a heterogeneous mixture of oligomers over biochemically-relevant concentration ranges.(10, 22, 23) We demonstrate the remarkable power of nMS to simultaneously measure liganded populations of different oligomeric states. Direct measurement of liganded populations has also revealed previously uncharacterized super-stoichiometric ATP binding to Rho oligomers. This capability can enable a rigorous treatment of the microscopic equilibria driving NTP turnover in hexameric helicases, with implications for characterizing thermodynamic properties in other complex biological milieux.

## Results and Discussion

Nano-electrospray ionization of ligand-free Rho produced a complex mass spectrum featuring a series of ions that could be assigned to the full range of oligomeric species expected for partially assembled hexameric protein rings, including pentamers, tetramers, trimers, dimers, and monomers (Figure 1B). The charge state distribution of each of these species is narrow and highly reproducible, and dissociation of the higher molecular weight species confirms the monomeric mass (Figure S1). However, we found that the relative abundance of the oligomeric species varied between experiments, independently of concentration or solution conditions, and were not a reliable metric of oligomeric population (Figure S2).

Heterogeneity in the oligomeric distribution could be confirmed by solution experiments. Size-exclusion chromatography (SEC) of Rho revealed that the centroid of the elution volume has a strong concentration dependence over a range from 0.1 µM to 10 µM Rho (Figure 1C). When loaded onto the analytical size-exclusion column at a protomer concentration of 10 µM, we observe a major peak eluting with a retention volume between that expected for a pentamer and hexamer (Figure S3). At 1 µM, the major peak broadens and shifts closer to the retention volume of the pentamer. At 0.1 µM, the major peak elutes between trimer and tetramer, and a second peak appears at the predicted retention volume of a monomer. Analytical SEC is limited by longitudinal diffusion contributing to already poor resolution between oligomeric states, and so we explored methods to measure oligomeric populations in solution at equilibrium.

Mass photometry (MP) experiments corroborated the existence of a heterogenous mixture of oligomeric species in solution. MP experiments conducted at 0.1 µM Rho revealed particle count distributions with molecular weights matching those expected of oligomeric states from monomer (47 kDa) through hexamer (283 kDa) (Figure 1D). The particle counts are highest for the monomer and hexamer at 0.1 µM. The lower particle counts of intermediate oligomeric states is suggestive of weakly positive cooperativity of self-assembly.(36, 37) MP experiments at 0.05 and 0.2 µM revealed a shift from distributions favoring monomer to hexamer (Figure S4), indicating an apparent dissociation equilibrium constant near 0.1 µM under our experimental conditions.

Because nMS enabled resolution of signals from each oligomeric state, we conducted titration experiments to simultaneously monitor ATP binding to each of the observed Rho oligomers. Importantly, upon optimizing experimental conditions to minimize salt/solvent adduct formation (Figure S5), we observed that – unlike the relative intensities between *different* oligomeric states – these conditions did not affect the relative intensities of signals from differently liganded states for a given oligomer. This allowed us to simultaneously and independently monitor ATP binding to each oligomeric state.

Spectra of 10 µM Rho equilibrated with equimolar ATP produced a resolved array of signals reflecting multiply charged ligand-bound states for each oligomeric species (Figure 2A). For each Rho oligomer, the mass shifts from the apo peak position are consistent with ATP binding (+507 Da, +530 Da if mono-sodiated). Observed masses under these “soft-ionization” conditions were slightly higher than the predicted sequence mass due to non-covalent adducts that were lost upon unfolding (Figure S1). The ratio of salt/solvent adducted to non-adducted species was insensitive to the presence of ATP, and so we conclude these adducts do not compete for ATP binding sites.

**Figure 2.**
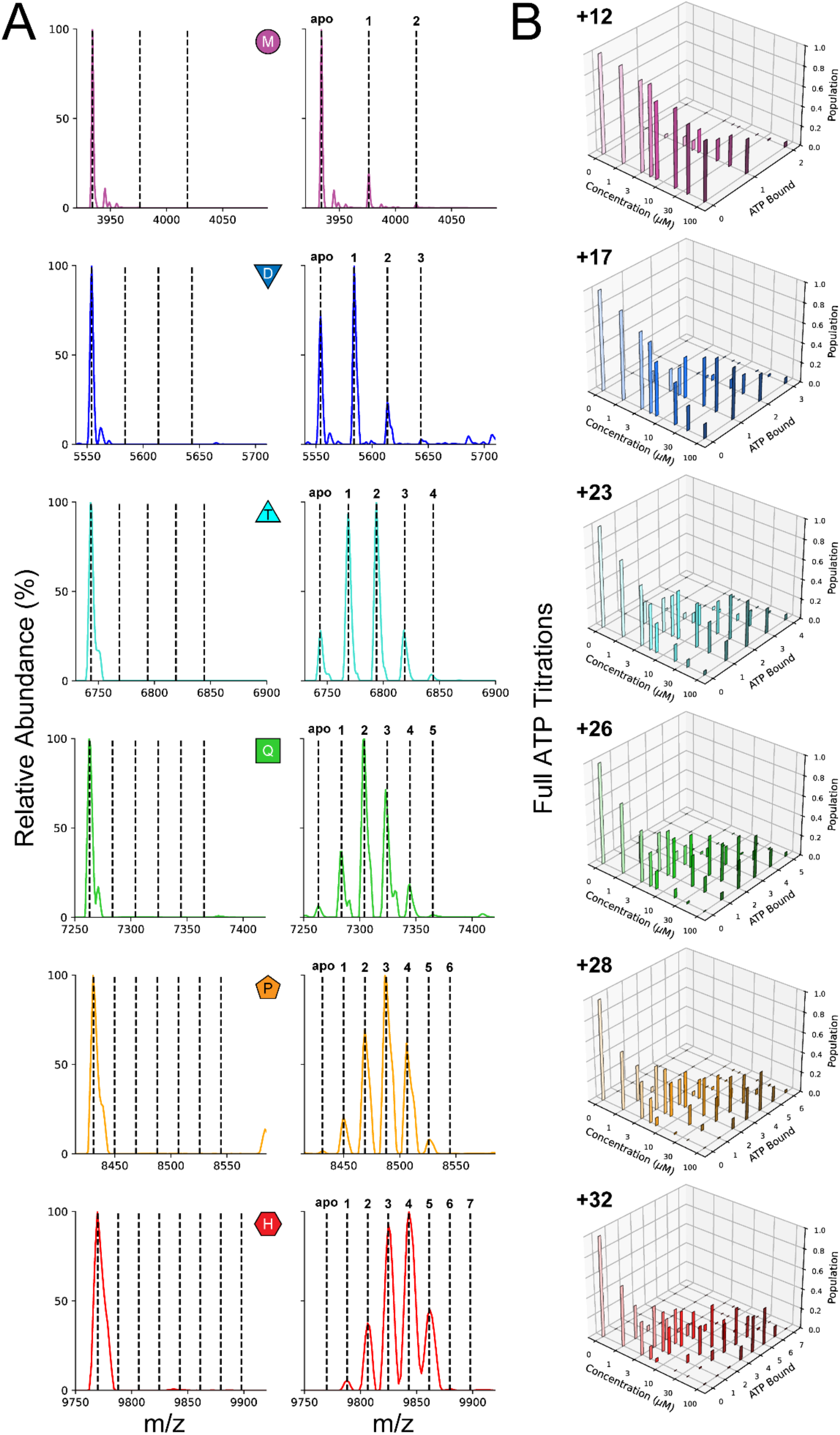
Native mass spectrometry enables simultaneous measurement of ATP-bound populations of each Rho oligomeric state. A) Representative mass spectra in the first two columns were collected with IST set to 100 V, a Rho protomer concentration of 10 µM, and ATP concentrations of 0 (left) and 10 µM (right). Dashed lines are drawn at the observed apo signal and predicted ATP-bound masses. The predicted peak positions for trimer through hexamer match to the mass of ATP plus 23 Da; sodiation is attributed to the two molar equivalents from the ATP salt. For the monomer through tetramer species, additional peaks are observed that correspond to super-stoichiometric binding at 10 µM ATP. B) Liganded populations from representative charge states over a range of ATP titrations were obtained from spectra recorded at IST set to 75 V; populations were obtained from the intensities of the signals. Super-stoichiometric binding to pentameric and hexameric Rho begins to emerge at 40 and 100 µM ATP, respectively.

An important observation is that the ligand binding pattern closely reflected the number of fully formed interfacial binding sites in each oligomeric species (Figure 1A, inset). For example, the dimer can properly assemble one enclosed ATP binding site and has two exposed, poorly formed sites. At 10 µM ATP, the dimer predominantly binds one ATP, with a smaller population binding a second ATP, and a trace population binding a third (Figure 2A). Likewise, for the trimer and tetramer, the binding pattern is consistent with well-formed sites binding efficiently, while species with one or two additional bound ligands are less abundant (Figure 2A). This pattern is also observed for the pentamer and hexamer at higher ATP concentrations (Figure 2B). Throughout the titration, site occupancy within each oligomeric state increased with rising ATP concentration (Figure 2B). However, none of the Rho oligomers reached full saturation over the concentrations sampled (up to 100 µM ATP).

These results may inform conflicting reports concerning the binding of ATP to Rho.(23, 38) Rather than reflecting weak binding to half of the sites in hexameric Rho, the observed multiphasic ATP binding behavior may instead arise from a population of partially formed binding sites. While interactions between sites could contribute to homotropic cooperativity, the presence of distinct classes of binding sites complicates interpretation of averaged isotherms. Such aggregation of complex binding events can obscure key features of the data – features which mass spectrometry is uniquely capable of capturing.

The power of nMS to resolve populations of liganded species, and their dependence on ligand concentration, has been demonstrated for quantifying cooperativity in ligand binding to ring-forming proteins.(2, 3) ATP binding to Rho monomers and dimers has been reported under less-native conditions,(12, 22) while the more complete and simultaneous characterization of ATP binding to trimer, tetramer, and pentamer are important advances of the current work. Identification of super-stoichiometric ATP binding has implications for prior work characterizing the two ATP binding modes of Rho. We anticipate that advances in measuring liganded populations of oligomeric states in the gas phase will enable full statistical thermodynamic modeling of ligand binding in Rho.(2, 5, 32) Such measurements may help us understand how ATP binding, hydrolysis, and exchange are coordinated to enable processive translocation along mRNA substrates to terminate transcription.

## Methods

### Recombinant Rho expression and purification

Electrocompetent T7 Express *lysY* cells (New England Biolabs) were transformed with a pET24b vector bearing the *E. coli* Rho gene modified with an N-terminal Met-Gly-His insert (Addgene catalog #113121; ProtParam predicted molecular weight post-N-Met processing: 47,198.40 Da; ε_280_ 15,930 M^-1^ cm^-1^) using a Bio-Rad MicroPulser to apply a 1.8 kV pulse across a 1-mm capacitor gap. Nascent transformants were spread on LB-agar plates infused with 50 mg/L kanamycin and 25 mg/L chloramphenicol and allowed to grow overnight at 37 °C. Isolated colonies were transferred to 50 mL starter cultures supplemented with 50 mg/L kanamycin and grown overnight. All liquid cultures were grown in a shaking incubator set to 37 °C and 220 rpm.

A 2 L flask holding 1 L of LB supplemented with 50 mg/L kanamycin was inoculated with 10 mL of the overnight starter culture. Protein expression was induced at an OD_600_ of 0.4 to 0.6 A using 1 mM isopropyl β-D-thiogalactopyranoside (IPTG). Recombinant protein expression proceeded for 3 hours under otherwise constant environmental conditions. The OD_600_ at harvest was typically between 1.5 and 1.8 A. Cells were separated from the liquid medium via centrifugation at approximately 4,200 RCF for 30 minutes in an SLA-3000 Super-Lite rotor (Sorvall). Cell pellets were immediately resolubilized in 30 mL of lysis buffer (Table S1; 50 mM Tris, 250 mM KCl, 1 mM TCEP, 10% glycerol at pH 7.6) supplemented with cOmplete Mini Protease Inhibitor Cocktail (MilliporeSigma).

The resolubilized cell slurry was lysed on ice using a Q125 Sonicator (QSonica) equipped with a 1/4” probe set to pulse for 5 seconds on, 10 seconds off at 50% amplitude for 5 minutes. The lysate was clarified of cellular debris via centrifugation at approximately 26,900 RCF for 30 minutes in an SS-34 rotor (Sorvall). Clarified lysate was passed through a 0.2 µm filter (Pall Corp.) prior to loading onto a pre-equilibrated 5 mL HiTrap Heparin HP column (Cytiva). Recombinant Rho was eluted from the heparin resin via a gradient from 0 to 1 M NaCl at 4 °C. Rho fractions were further purified via preparatory scale size-exclusion chromatography using a HiPrep 16/60 Sephacryl S-300 HR column (Cytiva) equilibrated in SEC buffer (Table S1; 20 mM Tris, 200 mM KCl, 0.2 EDTA, 0.2 mM DTT, 5% glycerol at pH 7.6) at 4 °C. Protein yields ranged between 30-50 mg as determined by A_280_.

### Analytical size-exclusion chromatography

Purified Rho was serially diluted to a protomer concentration of 10 µM, 1 µM, and 0.1 µM in 240 µL of SEC buffer and loaded onto a pre-equilibrated Superdex 200 Increase 10/300 GL column. The column was calibrated using Bio-Rad Gel Filtration Standard #1511901 reconstituted in Rho SEC buffer (Figure S3). All analytical SEC experiments were conducted in SEC buffer.

### Mass photometry data acquisition and processing

Static mass photometry measurements of purified Rho were collected on a Refeyn TwoMP. High-precision borosilicate microscope slides were batch-prepared with alternating cleanings of water and isopropanol before being dried with nitrogen gas and stored until use. Six-well gaskets were placed on the center of the prepared slides and mounted on the laser stage of the mass photometer. For each sample condition, a mass calibration was performed with a known calibration standard. In each well, to a maximum volume of 20 µL, the Rho stock was diluted to various concentrations in the low- to mid-nanomolar range using SEC buffer. Stage position, automatic focusing, and measurement recording were completed with AcquireMP software by Refeyn.

Using DiscoverMP by Refeyn, the ratiometric contrast recordings were processed into a histogram of ratiometric contrast to scattering signal. By applying a mass standard calibration, Gaussian fitted peaks were assigned masses and Rho oligomers were identified.

### Sample preparation for native mass spectrometry

Purified Rho was exchanged from storage buffer into 100 mM EDDA (Table S1, CAS No. 5657-17-0) using an Amicon Ultra centrifugal filter with a MWCO of 10,000 Da. The filter was centrifuged at approximately 13,400 RCF until the volume in the well dropped from its maximum to half capacity (approximately 200 µL). The well was then refilled and this process repeated until the concentration of KCl dropped below one-sixth of the protomer concentration.

Disodium ATP salt (CAS No. 51963-61-2) was dissolved in EDDA and pH adjusted to 7.1 using ammonium hydroxide (CAS No. 1336-21-6) and acetic acid (CAS No. 64-19-7). The ligand solutions were diluted from concentrated stocks under buffered conditions.

### Native mass spectrometry data acquisition

Native mass spectrometry datasets were acquired using a Thermo Q Exactive UHMR Hybrid Quadrupole-Orbitrap with an in-house SID device in place of the transport multipole.(39, 40) Capillary emitters were prepared using a P97 micropipette puller (Sutter Instrument Company).(41) Emitters were mounted onto a Nanospray Flex Ion Source modified with platinum wire. CsI clusters were utilized as a monoisotopic calibration standard in positive ion mode. Spectra of unfolded monomers were obtained via aggressive activation.

Protein-ligand mixtures were prepared at room temperature in microcentrifuge tubes before loading into capillary emitters. IST voltages were arrayed at each titration point. 25 to 50 scans were collected per dataset, or approximately 30 seconds to 1 minute of data acquisition.

### Native mass spectrometry data processing and analysis

Xcalibur (Thermo) was used to access data in RAW format. Intensities averaged over the acquisition period were extracted as text files and ordered according to their mass-to-charge ratio. Automated deconvolution via UniDec was confirmed by manual assignment of charge states.(25) Local maxima were extracted using UniDec’s Data Collector module configured to the deconvolved apoenzyme mass plus zero to seven bound ATP molecules.(42) A window size of 2 m/z was utilized to account for variation in peak position due to adducting species.

Liganded populations were calculated by normalizing the local maxima by the sum of the extracted intensities. Titrations were performed at a protomer concentration of 10 µM.

## Acknowledgements

This work was supported by the Native MS-guided Structural Biology Center (NIH grant 1RM1GM149374 to VW), the Campus Chemical Instrument Center (OSU), and a seed grant from the College of Arts and Sciences (OSU). We thank Irina Artsimovitch (OSU) for reagents and advice, and the Foster and Wysocki research group members and James Berger (JHU) for encouragement and stimulating discussions.

