## Supporting Information for "Simultaneous ligand binding to intact and partially formed ATP binding sites in the hexameric termination factor Rho"

**Corresponding Author**

*Mark P. Foster

Mailing address: 734 Riffe Building, 496 West 12th Avenue, Columbus, Ohio 43210

Landline: 614-292-1377

**Author Contributions**

**Tyler D Billings**: Writing, Conceptualization, Resources, Methodology, Formal Analysis, Visualization.

**Kristie Baker**: Writing, Investigation.

**Philip Lacey**: Writing, Investigation, Formal Analysis, Visualization.

**Rodrigo Muzquiz**: Conceptualization, Investigation, Methodology.

**Vicki H. Wysocki**: Writing, Resources, Supervision, Funding Acquisition.

**Mark P. Foster**: Writing, Conceptualization, Formal Analysis, Supervision, Funding Acquisition.

Table S1. Buffers used to prepare and store Rho and its ligands. FPLC buffers were degassed under vacuum.

| Buffer Name | Component Concentration | Recipe |
|---|---|---|
| **Storage/SEC** | 20 mM Tris at pH 7.6<br>200 mM KCl<br>0.2 mM EDTA<br>0.2 mM DTT<br>5% glycerol* | • 2.341 Tris HCl, 0.624 g Tris Base<br>• 14.91 g KCl<br>• 0.058 g EDTA<br>• 0.031 g DTT<br>• 50 g glycerol |
| **Lysis** | 50 mM Tris at pH 7.6<br>250 mM KCl<br>1 mM TCEP<br>10% glycerol | • 6.233 g Tris HCl, 1.266 g Tris Base<br>• 18.64 g KCl<br>• 0.250 g TCEP<br>• 100 g glycerol |
| **Heparin A** | 10 mM Tris at pH 7.6<br>0.1 mM EDTA<br>0.1 mM DTT<br>5% glycerol | • 1.17g Tris HCl, 0.312 g Tris Base<br>• 0.029 g EDTA<br>• 0.015 g DTT<br>• 50 g glycerol |
| **Heparin B** | 10 mM Tris at pH 7.6<br>1 M NaCl<br>0.1 mM EDTA<br>0.1 mM DTT<br>5% glycerol | • 1.17 g Tris HCl, 0.312 g Tris Base<br>• 58.44 g NaCl<br>• 0.029 g EDTA<br>• 0.015 g DTT<br>• 50 g glycerol |
| **ESI Buffer** | 100 mM EDDA at pH 7.1 | • 17.617 g EDDA |

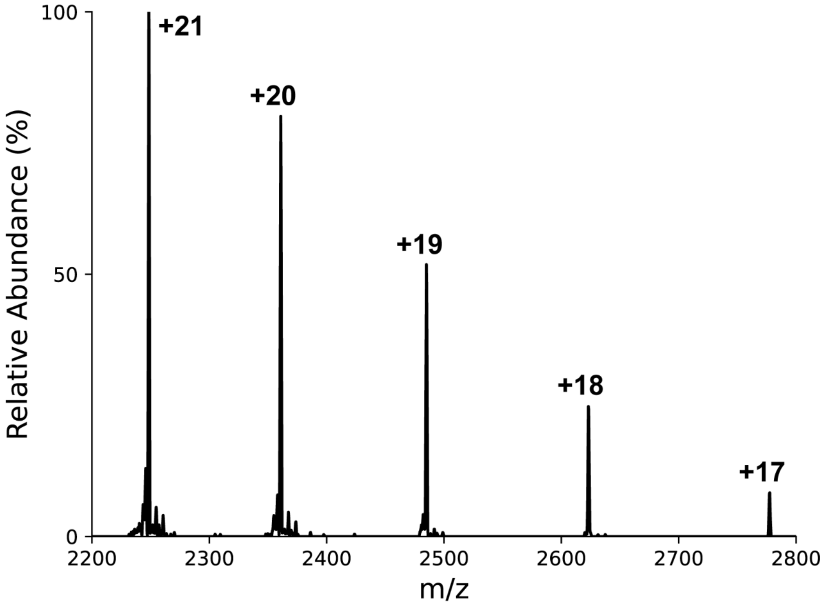


*Figure S 1. Intact monomer mass is confirmed from multiple charge states in a mass spectrum generated by activating 2 μM Rho in 300 mM ammonium acetate. Spectrum was recorded using a Thermo Q Exactive UHMR Hybrid Quadrupole-Orbitrap with an in-house SID device in place of the transport multipole.(1, 2) Rho was activated using IST 200 and HCD 200. The high charge is indicative of an unfolded monomer. The peak positions are as follows: 2777.41 (+17), 2623.09 (+18), 2485.03 (+19), 2360.87 (+20), and 2248.49 (+21). The deconvolved mass from these peaks is 47,197.57 ± 0.87 Da. The expected molecular mass post-N-Met processing is 47,198.40 Da. The predicted mass is slightly larger than Rho as found in E. coli strain K-12 (UniProt accession P0AG30, 46,873.02 Da) due to an N-terminal expression tag (Met-Gly-His-EcoRho) incorporated by the Berger group.(3)*

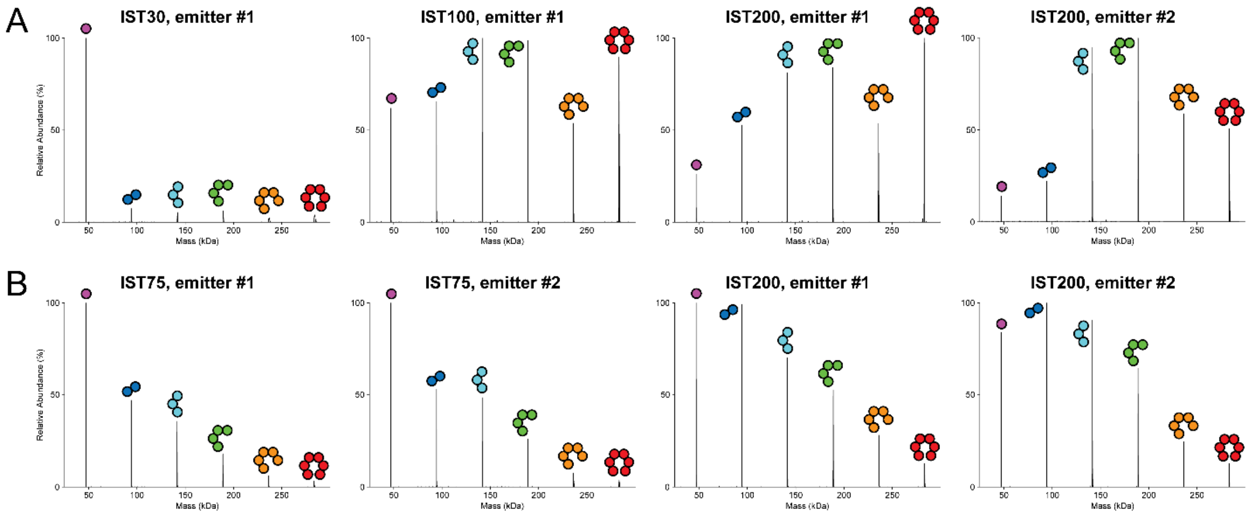


*Figure S 2. Signal intensity for different oligomeric states of Rho in native mass spectra was highly variable, independent of concentration. Top row (A) is 10 µM Rho, bottom row (B) is 2 µM Rho. In-source trapping (IST) voltages and emitter ID are labeled for each plot. Populations are derived from UniDec deconvolution(4), with mass range of 25,000 Da to 300,000 Da, automatic m/z peak width determination turned off, Softmax smoothing of 10 and point smooth width of 10. Relative intensities are observed to vary with activation and emitter (capillary tip), making these spectra poor metrics of the population of each oligomeric state.*

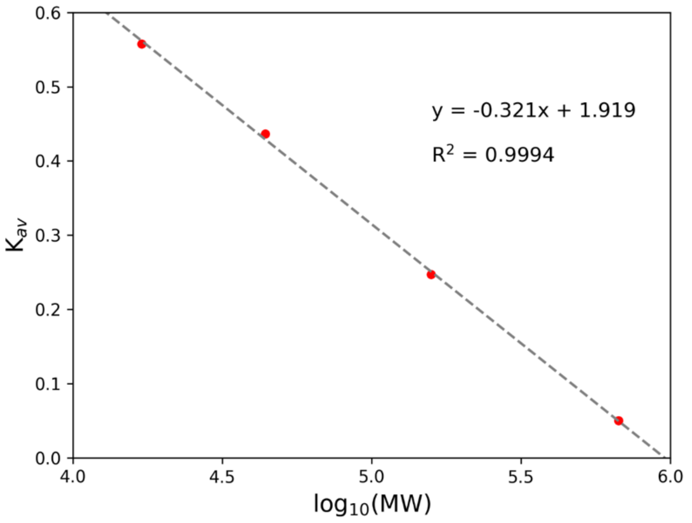


*Figure S 3. Calibration curve to predict elution volumes of varying EcoRho oligomeric assemblies. Protein standards are bovine thyroglobulin (670,000 Da), bovine gamma-globulin (158,000 Da), chicken ovalbumin (44,000 Da), and horse myoglobin (17,000 Da). Vitamin B12 (1,350 Da) was excluded from the calibration as its molecular weight lies outside the linear range of the column.* $K_{av}$ *is determined from:*

$$K_{AV} = \frac{(V_e - V_0)}{(V_t - V_0)}$$

*where $V_e$ is the specific elution volume of the analyte, $V_0$ is the void volume of the column, and $V_t$ is the total volume of the column. The regression statistics are reported in the plot.*

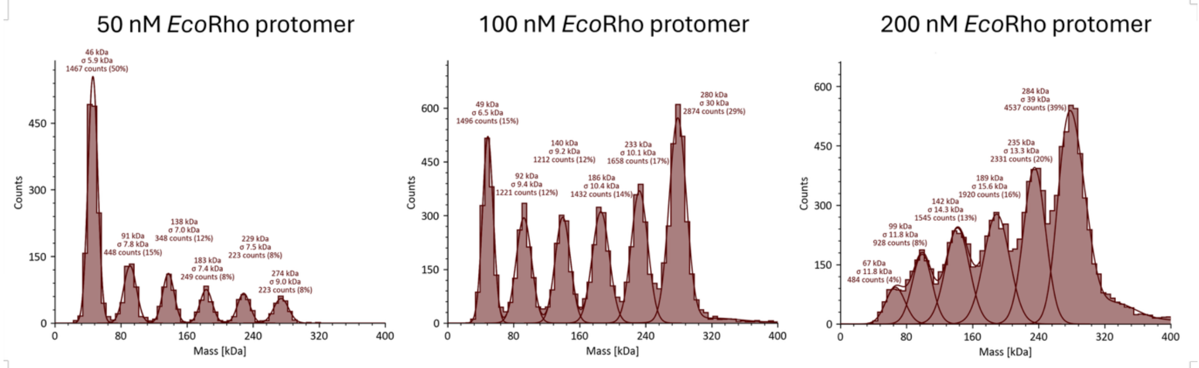


*Figure S 4. E. coli Rho oligomerization can be measured by mass photometry. The 100 nM measurement is a replicate collected on a different day from the data presented in Figure 1D. The relative abundance of oligomeric species is suggestive of positively cooperative self-assembly. Protomer concentrations higher than 200 nM were also assayed, but the particle count distributions coalesce to an average molecular weight of 400 ± 100 kDa.*

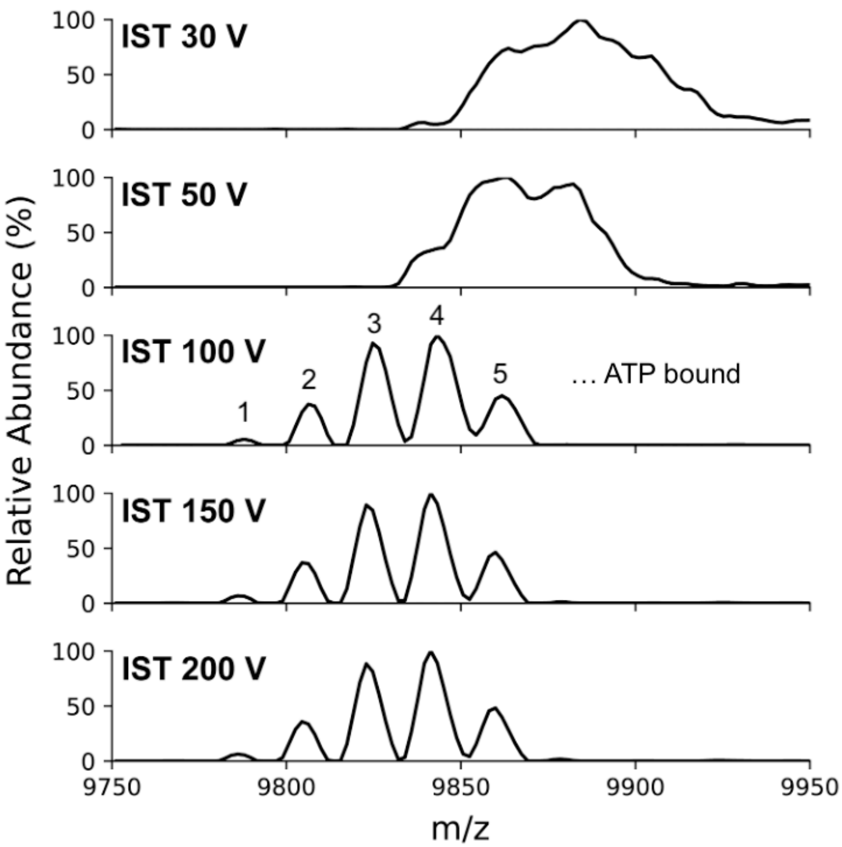


*Figure S 5. Moderate in-source trapping does not alter relative intensities of differently ATP-bound Rho. The +29 charge state of the hexamer is used as an example. The peaks in the 100 V panel were assigned to the indicated ATP-bound states shown in Figure 2. 75 V was selected for titration experiments as sub-50 V resulted in poor resolution.*
